# Assessing contamination in DNA extraction kits commonly used for microbiome research

**DOI:** 10.1101/2025.09.23.677068

**Authors:** Jorden T. Rabasco, Samantha C. Kisthardt, Casey M. Theriot, Benjamin J. Callahan

## Abstract

Sequencing-based measurements are now routinely used to investigate the microbial world, however, contamination by DNA outside the intended sample remains a problem. Contaminants obscure the true microbial signal and can lead to misleading scientific interpretations. Much work has been done to address the effects of these contaminants including best practices as outlined in *Eisenhofer et al*. and *Fierer et al*. Yet, even with best practices in place, the current literature consensus is that contaminants remain impactful, at least in low biomass environments (5, 7, 11, 13, 16). One well-known source of contaminants are those found within DNA extraction kits, as was shown clearly in the pioneering work of *Salter et al. 2012* and *Karstens et al. 2019*. However, given the rapid evolution of DNA sequencing methods, it would be worthwhile to revisit the issue of contaminants in contemporary DNA extraction kits (the “kitome”). Here we provide an updated characterization of the ‘kitomes’ of DNA extraction kits commonly used for microbiome research.

**Importance:** Microbial contamination in commonly used DNA extraction kits has not been recently assessed. Here we evaluate the contamination in DNA extraction kits commonly used in microbiome studies over the past several years, and provide actionable guidance on appropriate DNA extraction kits for low biomass microbiome measurements.

## Body

Sequencing-based measurements of microbial communities have given us a new lens into the microbial world that is free from some previous constraints, such as culturability (4). However, the accuracy of sequencing-based measurements can be degraded through mechanisms such as sequencing error, amplification bias, and of particular interest to us here – contamination. Contaminants are defined as any DNA molecule or read in the output sequencing file not originating from the community of interest. Contaminants can come from a variety of sources including the sampling environment (3, 17), the microbiome within the lab space (10), and the kits and reagents used in the experimental process (11, 12, 16). DNA extraction kits have been shown to be an important contributor to overall contamination in the past, with their contribution to contamination in microbiome measurements earning the appellation of the ‘kitome’ (14). Measurement errors caused by contamination can lead to false positive or negative results (14, 16), altered relative abundances (3, 16) and skewed diversity measurements (6). Here, we revisit the ‘kitome’ in the current landscape of widely used DNA extraction kits in microbiome research, and provide actionable guidance on choice of DNA extraction kits for low-biomass microbiome measurements.

We identified eight DNA extraction kits (Table 1) that are popularly used in microbiome papers employing sequencing-based measurements (see Methods). We procured examples of all eight of these kits in 2024. For three of these kits – Zymo Biomics Quick-DNA Minprep Kit, Macherey-Nagel NucleoSpin Triprep, and the Qiagen DNeasy Blood and Tissue kit – we were also able to obtain a version of the kit procured at least four years earlier, allowing an evaluation of potential change over time in their ‘kitome’.

**Table 1.** DNA extraction kits included in this study.

| Kit | Manufacturer | Year Procured |
| --- | --- | --- |
| DNeasy Blood and Tissue Kit | Qiagen | 2024 |
| DNeasy Blood and Tissue Kit | Qiagen | 2020 |
| DNA miniprep kit | Zymo | 2024 |
| Quick-DNA Miniprep kit | Zymo | 2024 |
| Quick-DNA Miniprep kit | Zymo | 2020 |
| DNeasy PowerSoil Pro DNA Isolation Kit | Qiagen | 2024 |
| Nuclospin TriPrep | Macherey-Nagel | 2024 |
| Nuclospin TriPrep | Macherey-Nagel | 2015 |
| QIAamp Fast DNA Stool Mini Kit | Qiagen | 2024 |
| QIAamp DNA Microbiome kit | Qiagen | 2024 |
| QIAamp Powerfecal Pro Kit | Qiagen | 2024 |

We developed a dilution series of a bacterial monoculture in order to evaluate contamination at varying concentrations of input bacteria, following the examples of *Salter et al*. and *Karstens et al*. Specifically, the stbl04 strain of *E. coli* was grown overnight in LB broth, from which aliquots were pelleted and frozen (Methods). The concentration of the initial *E. coli* monoculture was quantified by plating. For each kit, three pellets were thawed and reconstituted in 222ul of PBS, and then a five-step 10x dilution series (D0 - D5) was generated. The undiluted D0 sample contained ∼2.7×10^7^ CFUs in 200uL, and the most diluted D5 samples contained ∼270 CFUs in 200uL. All extractions took place in sterile conditions and followed the manufacturer’s instructions. After DNA extraction was complete, all samples were eluted in a 50ul volume either using the provided elution buffer if included in the kit or molecular grade water, (sterile, filtered) if not.

Extracted DNA concentrations varied across kits. Most kits yielded ∼1-4 ng/ul. However, the DNeasy Blood and Tissue Kit and both Nucleospin Triprep Kits yielded lower concentrations in the ∼0.1-0.8 ng/ul range, while the QIAamp Fast DNA Stool Mini Kit yielded DNA concentrations too low to quantify with the high-sensitivity Qubit protocol. We performed DNA quantitation on the full range of dilution samples, and in all kits saw the expected behavior of DNA concentrations falling by approximately 1/10th at each step in the 1:10 dilution series.

We measured the composition of each sample by short-read 16S rRNA gene sequencing (Methods). After denoising, amplicon sequence variants (ASVs) from each sample were assigned as ‘expected’ if they exactly matched the known 16S rRNA gene sequence of *E. coli* Sbtl04, and ‘contaminant’ if they did not. While contaminants did rise in abundance with increased dilution as expected, in most kits the fraction of contaminant ASVs remained low at moderate and even high dilutions – typically <4% contaminants at D3 (∼2.7×10^4^ CFUs), apart from two replicate outliers, and typically <20% contaminants at D5 (∼2.7×10^2^ CFUs). 5 of the 8 kit types tested conformed to this pattern which is contrary to past experimentation. In *Salter et al*. their dilution series, denoted as Sal_D0-Sal_D5, showed their most dilute samples (2×10^3^ CFUs), as having ∼80% contamination, and were on the same dilution scale as our D4 samples (2.7×10^3^ CFUs), which showed typically <10% contamination. *Karstens et al*.’s dilution series, denoted as Kar_D0-Kar_D8, was much more concentrated than ours with a starting concentration of ∼1.5×10^9^ CFUs/sample, compared with our starting concentration of ∼2.7×10^7^ CFUs/sample. The dilution series utilized in *Karstens et al*. displayed substantially more contamination across their samples, with their Kar_D6 samples (1.44×10^6^ CFUs) showing ∼65% contamination being comparable to our D1 samples (2.7×10^6^ CFUs) which showed ∼0.01% contaminants detected. Overall, contaminant ASVs were severely reduced in most kits across the dilution series when compared with previous experimentation (11, 16) barring the most dilute samples (Figure 1).

**Figure 1.**
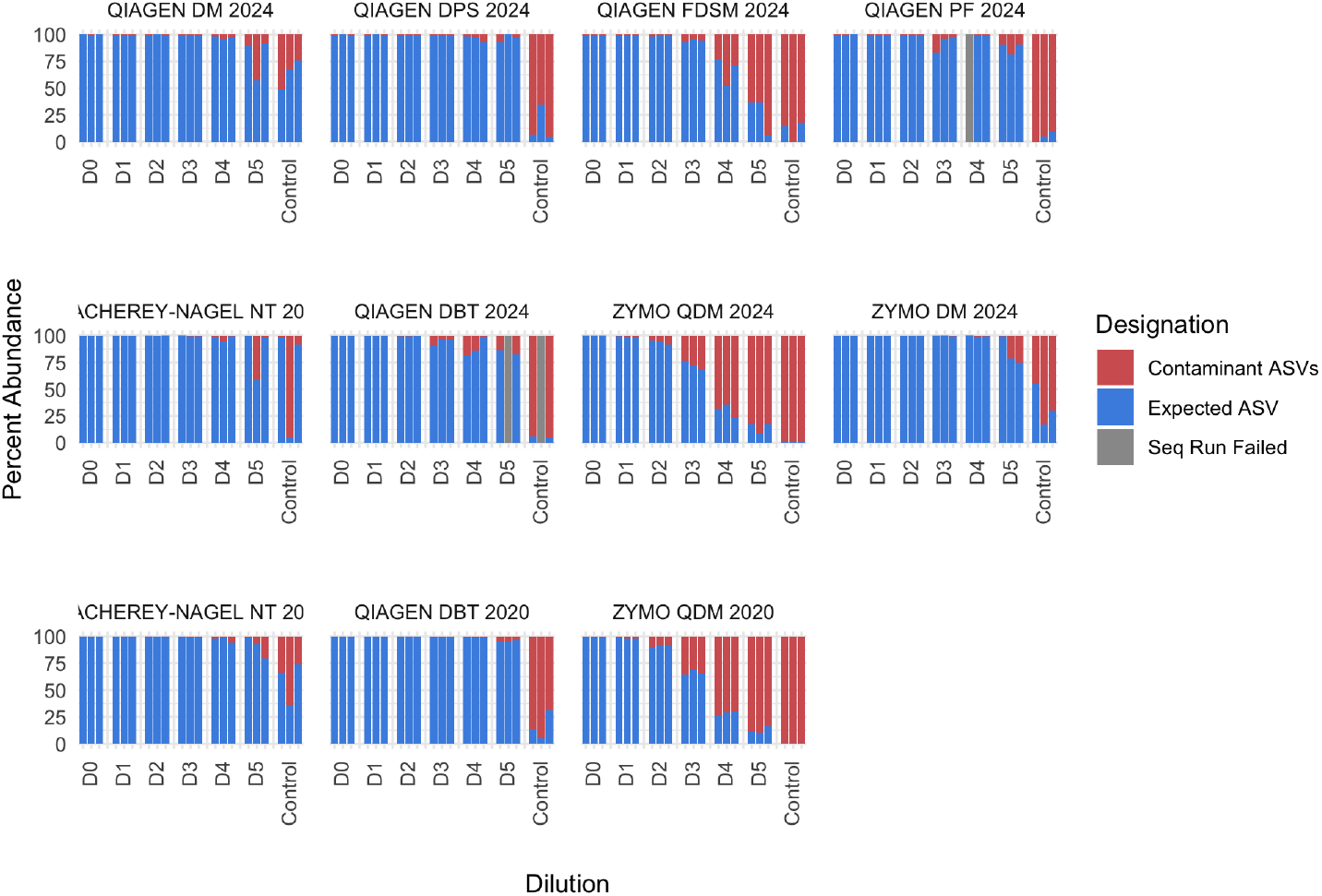
Relative abundance of expected ASVs and contaminants across a ten-fold dilution series by DNA extraction kit. D0: ∼1.35×10^8^ CFUs/mL; D1: ∼1.35×10^7^ CFUs/mL; D2: ∼1.35×10^6^ CFUs/mL; D3: ∼1.35×10^5^ CFUs/mL; D4: ∼1.35×10^4^ CFUs/mL; D5: ∼1.35×10^3^ CFUs/mL; Control: No input cells.

We observed a higher level of contamination in three kits relative to the rest – Zymo Quick DNA Miniprep Kit (2020, 2024) and Qiagen QIAamp Fast DNA Stool Mini Kit (2024). In these three higher-contamination kits, over 25% of sequencing reads were assigned to contaminants by the 4th dilution (1:10,000). The higher level of contamination associated with the QIAamp Fast DNA Stool Kit may be explained in part by the low extraction efficiency of that kit in our experiment: extracted DNA concentrations from this kit were too low to quantify even in the undiluted sample. The two examples of the Zymo Quick DNA Miniprep Kit (2020 and 2024) showed nearly identical patterns of increasing contamination in more dilute samples, with both yielding >25% contaminants by the third dilution and >60% contaminants by the fourth dilution.

The taxonomic identity of contaminant ASVs in all samples were assessed (Figure 2). In the kits without significant contamination those contaminant ASVs detected at high dilutions or in the reagent blanks were taxonomically diverse and often could not even be assigned to the genus level, perhaps reflecting off-target amplification. The Zymo Quick DNA Miniprep Kits had a unique contaminant profile consisting almost entirely of a single *Pseudomonas* ASV. This consistency is particularly notable given the four year time gap between the procurement of the 2020 and 2024 examples of this kit.

**Figure 2.**
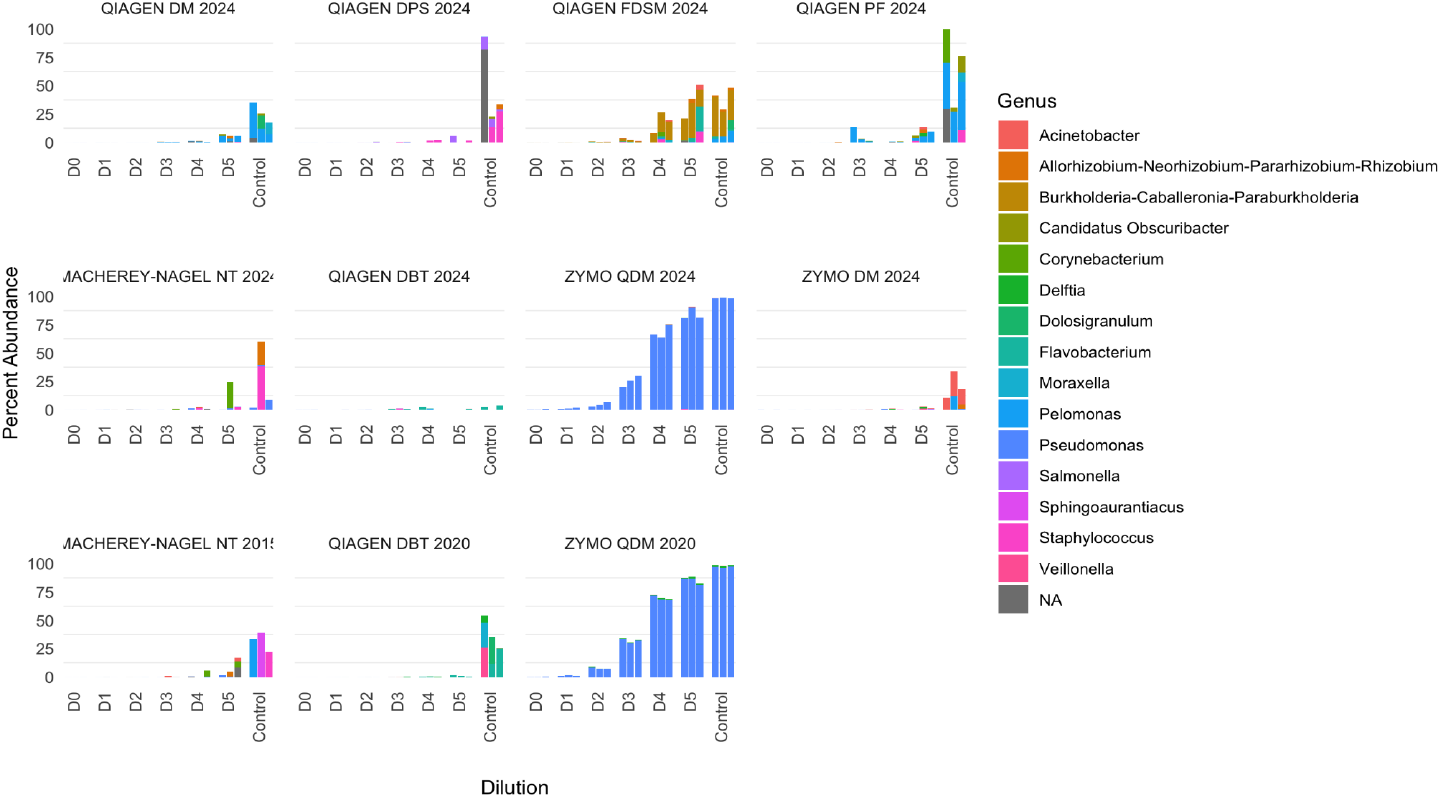
Taxonomic assignment of contaminants as a function of dilution and DNA extraction kit.

## Discussion

Two previous landmark papers investigating the “kitome” in low biomass microbiome sequencing – *Karstens et al*. and *Salter et al*. – reported a strong correlation between lower input biomass and higher contamination, with contaminants making up a substantial (>80%) fraction of sequencing reads for samples with ∼2×10^3^ CFUs. In contrast, here we found that – for a majority of the kits tested – contamination remained <10% in even very dilute samples with ∼2.7×10^3^ CFUs. All three studies used 16S rRNA gene sequencing of a dilution series of a simple input (monoculture or mock community) to assess contamination as a function of input cell count. Despite modest differences in experimental specifics (e.g. PCR cycle count), we believe results can be directly compared across these studies as a function of input cell number, which spanned a range of 2.7×10^7^ - 2.7×10^2^ CFUs/sample in our study versus 2×10^8^ - 2×10^3^ CFUs/sample in *Salter et al*. and 1.05×10^9^ - 1.6×10^5^ CFUs/sample in *Karstens et al*. (determined from the Zymo mock community product specification). This comparison shows that most DNA extraction kits in common current usage have meaningfully lower contaminant burdens than the older kits evaluated in those previous papers.

Our results provide straightforward guidance on the choice of DNA extraction kit for low biomass microbiome sequencing. Most clearly, we recommend against using the Zymo Quick DNA Miniprep Kit or the QIAamp Fast DNA Stool Kit for samples with <2.7×10^4^ CFUs, while the other six kits tested get a “clean bill of health” for even very dilute samples. In addition, if *Pseudomonas* is an expected member of sampled microbial communities, then the Zymo Quick DNA Miniprep Kit should probably be avoided entirely. That said, we also want to emphasize the apparently high production consistency of the Zymo kit – the exact same *Pseudomonas* ASV in the same quantities was present in kits procured in 2020 and 2024. Furthermore, investigators should heed manufacturer guidance when choosing reagents. The Zymo DNA Miniprep Kit (no “Quick”) is specifically designed for microbiome applications as reported by the manufacturer, and was found to be essentially free of contaminating microbial DNA.

Overall, we believe our results show that multiple DNA extraction kits are now available that are appropriate for low biomass microbial sequencing. This may reflect increased attention from manufacturers on reducing microbial contamination in their products since the rise in importance of microbial sequencing over the past 15 years. We also caution that although we report relatively low levels of contaminants from most DNA extraction kits tested, contamination can arise from many steps in the sampling and measurement process. Care and attention to all aspects of this workflow is necessary to control contamination in low biomass sequencing, see the recent consensus statement for more comprehensive guidance (9).

## Methods

### Selection of DNA Extraction Kits

We performed a Google Scholar search using the terms “microbiome” and “DNA extraction”, restricting results to papers published since 2020. We tallied the number of times each DNA extraction kit appeared in the methods section of the top 50 papers returned by this search. We selected the five kits that appeared in the most papers to be included in our study: Qiagen PowerSoil DNA Isolation Kit, QIAamp Fast DNA Stool Mini Kit, QIaamp DNA Microbiome kit, QIAamp PowerFecal Pro DNA Kit, ZymoBIOMICS DNA Miniprep Kit.

We augmented this set of kits with three more kits for which we were able to obtain complete older (2020 or earlier) examples of the kit: Qiagen DNeasy Blood and Tissue Kit, Macherey-Nagel NucleoSpin Triprep Kit, ZymoBIOMICS Quick-DNA Minprep Kit. Having paired old and new examples of these kits allowed us to test if contamination profiles in these kits had changed over time. Of note, DNeasy Blood and Tissue Kit is the same kit used in the *Karsten et al*.*’s* study on which our experimental design is partially modelled. In sum, we included in our study eleven example kits representing eight DNA extraction kit products.

### E.coli Pellet Generation

An overnight culture of *Escherichia coli* Sbtl04 was prepared by inoculating 5 mL Sigma-Aldrich LB (Lennox) broth (catalog #L3022) with a single colony of *E*.*coli* and incubating at 37°C overnight, shaking at 225 RPM. The next day, the overnight OD was measured, and a 10 mL culture was prepared at a starting OD of 0.1. This culture was allowed to grow until it reached an OD of ∼0.6, and plated to assess a quantification of 2×10^8^ CFU/ML. The culture was then split into 40 aliquots of 150 μL each. The aliquots were centrifuged at 4900xg for 3 minutes and the supernatant was aspirated. Immediately following removal of the supernatant, the pellets were frozen at -80°C.

### Dilution Series Generation

Undiluted samples were created by reconstituting one *E. coli* pellet (approximately 3×10^7^ CFUs) in 222uL of Phosphate Buffer Solution (PBS). A 1:10 dilution was then created by transferring 22ul from the starting sample into 200ul of PBS. This procedure was repeated four more times, resulting in a dilution series spanning five orders of magnitude and where the most concentrated samples had ∼2.7×10^7^ CFUs and the least concentrated samples had ∼270 CFUs. Three independent replicate dilution series were generated for each DNA extraction kit.

### DNA Extraction and Sequencing

All DNA extractions were performed in a sterile class 2 Nuaire Biosafety Lab Cabinet. All centrifuges, shakers, and thermocyclers used in the DNA extraction process were first cleaned with Thermofischer RNase away (catalog #7002), followed by 70% ethanol. Dilution series samples were eluted in 50ul of the appropriate elution buffer for each kit prewarmed to 30C. Extracted DNA was quantified via a Qubit system and kept at -20C for sequencing by the Duke Microbiome Sequencing Core. The samples were sequenced on an Illumina Miseq (2x250bp) targeting the V4 region of the 16S rRNA gene after undergoing PCR with 35 cycles.

### ASV generation and Analysis

Amplion sequence variants (ASVs) were resolved using the software DADA2 (2), and taxonomically assigned with the Silva nr99 16S database (15) formatted for DADA2. ASVs were identified as ‘Expected’ if they exactly matched the V4 region of the 16S rRNA gene in the stbl04 *E. Coli* genome (CP076044) (1). All other ASVs were identified as contaminants.

## Acknowledgements

We thank the Duke University School of Medicine for the use of the Microbiome Core Facility, which provided library preparation and sequencing services. We acknowledge the Breen Lab, the Yoder lab, and the Sheahan Lab for assistance in sourcing DNA extraction kits procured prior to 2021. This work was supported primarily by the Engineering Research Centers Program of the National Science Foundation under NSF Cooperative Agreement No. EEC-2133504.

## Author Contributions

Conceptualization, JTR BJC and CMT; methodology, JTR BJC SCK and CMT, software, JTR and BJC; validation, JTR and BJC; formal analysis, JTR; investigation, JTR BJC SCK; resources, BJC and CMT; data curation, JTR and SCK; writing—original draft preparation, JTR and BJC; writing—review and editing, JTR BJC SCK and CMT; visualization, JTR; supervision, BJC; project administration, BJC; funding acquisition, BJC. All authors have read and agreed to the published version of the manuscript.

